# Cerebral amyloid angiopathy and comparative analysis of amyloid-β protein in birds

**DOI:** 10.1101/486340

**Authors:** Mutsumi Yamazaki, Maho Morimoto, Fuyuki Kametani, Steven James Scott, Dennilyn Parker, Anjana Chamal Karawita, Yumi Une

## Abstract

Senile plaques and cerebral amyloid angiopathy (CAA) are well-documented in various mammals, and several species even exhibit neurofibrillary tangle (NFT). However, we know far less about whether such symptoms are present in birds. Therefore, here we clarified the occurrence and pathogenesis of avian aβ-related lesions, analyzing the aβ amino-acid sequence across 28 birds at multiple life stages, representing 15 species, 14 genera, and 9 nine families. We also determined the expected aβ amino-acid sequence after comparing data from the brains of nine birds (seven species) with publicly available NCBI data. We observed CAA and senile plaque-like deposition only in a female Amazon parrot, estimated to be around 30–40 years old. We identified two Aβ depositions (40 and 42) in the same location that correspond to Aβ 6-42. Additionally, we observed severe Aβ deposition, accompanied by severe hemorrhaging, in blood vessels of the superficial and deep portions of the brain. These lesions were directly related to the cause of death. Of 40 bird species, 36 exhibited type 1 Aβ amino-acid sequences, similar to humans. Given that all of these birds were old, our results suggest that Aβ is deposited primarily as CAA as the animals age. This report is the first clinically based description of CAA in birds. Interspecific variation likely exists because we identified species that did not exhibit Aβ deposition even when the birds are old enough. However, even birds of the same taxonomic status differed in whether they possessed or lacked Aβ deposition. Thus, other factors besides Aβ amino-acid sequence could influence this symptom.

## Introduction

Amyloid β protein (Aβ, 40–43 amino acids) is produced when β and γ secretases cleave the N- and C-termini of the amyloid precursor protein (APP), respectively [1], but not when a secretase cleaves App [2]. The deposition of Aβ in human brain parenchyma and cerebral blood vessels induces dementia and cerebrovascular disorders. One particularly well-known neurodegenerative disorder of this kind is Alzheimer’s disease. Its pathological characteristics include three distinct lesions: senile plaques with aggregated Aβ, argyrophilic fibrous structures in neurons (neurofibrillary tangles, NFT), and generalized brain atrophy associated with neuronal loss [3]. The well-supported amyloid cascade hypothesis of Alzheimer’s disease suggests that the first pathological event is Aβ deposition, eventually leading to neuron death [4]. Cerebral amyloid angiopathy (CAA) refers to Aβ deposits in cortical and meningeal blood vessels. This condition occurs in Alzheimer’s patients and also frequently in the elderly [5]. Besides being reported in humans, CAA and other forms of Aβ deposition have been confirmed in numerous other mammals, including non-human primates, canines, felines, bears, marine mammals (sea lion), ungulates (Bactrian camel), and rodents (degu) [6]. In contrast, Aβ deposition has only been observed in three avian species: great spotted woodpecker (*Picoides major*) [7], Steller’s sea eagle (*Haliaeetus pelagicus*), and White eagle (*H. albicilla*) [8]. Owing to this lack of research, we know very little about interspecific differences among avian species in terms of the incidence of Aβ deposition. Furthermore, the actual mechanisms of this symptom remain unclear. Therefore, this study aimed to determine the pathological characteristics of Aβ-related brain lesions in birds. Additionally, we analyzed Aβ amino-acid sequences from multiple species to elucidate pathogenesis.

## Materials and Methods

### Study design

#### 1. Animals

Pathological appraisal was performed on 28 birds. Of these, 27 died of natural causes at multiple zoos in Japan (Hamura Zoo; Tokyo, Nagasaki Penguin Aquarium; Nanasaki, Kakegawa kachouen; Shizuoka, Fuji kacyouen; Shizuoka, Fukuoka City Zoological Garden; Fukuoka and Wild bird park; Saitama) and in two private collections. An amazon parrot was euthanized with sodium pentobarbital, 0.44ml/kg body weight, intraosseous (Ethanol Pentobarbital Sodium Injection, 240mg/ml, Bimeda-MTC Animal Health Inc., Cambridge, ON, N3C2W4) due to its poor clinical prognosis at the Western College of Veterinary Medicine. Analyses were performed in the Laboratory of Veterinary Pathology at Azabu University. The 28 subjects represented 15 species, 13 genera, and nine families (Table 1). Because average life expectancy varies across species, standardized life stage levels (LSL) ranging from young (1) to oldest-old (6) birds were determined (Table 1). The brains of eight birds, each representing a different species, were used to analyze Aβ amino-acid sequences. Publicly available NCBI data on an additional 40 birds (spanning 39 families) was added to improve interspecific comparisons (Table 2).

#### 2. Histopathological examination

The whole brain was fixed with 10% neutral buffered formalin solution. Paraffin blocks were made from the frontal section of the cerebrum (visual cortex), as well as sagittal sections of the cerebellum and spinal cord, and cut into 2 μm, 5 μm, and 7 μm. Hematoxylin-eosin, Kluver-Barrera, Holzer, Congo red, and periodic acid silver methenamine staining were performed to confirm the presence and morphology of senile plaques.

#### 3. Immunohistochemical examination

The following antibodies were used to evaluate Aβ deposition and morphology in brain tissues: Aβ 17-24 mouse monoclonal (4G8, Covance, NJ, USA) for detecting C-terminal amino acid, anti-human Aβ1-40 rabbit polyclonal (IBL, Gunma, Japan), and anti-human Aβ1-42 rabbit polyclonal (IBL, Gunma, Japan). Furthermore, hyperphosphorylated tau presence was confirmed using anti-human PHF-Tau and phosphorylated Ser202 antibody AT8 (INNOGNETICS, Gent, Belgium). For immunogen, dilution concentration, and activation treatment per antibody, see Table 3. The positive control for immunostaining was APP transgenic mice.

#### 4. Amino acid sequence analysis of avian Aβ

Using the brains of nine birds (representing seven species; Table 2), the APP gene was sequenced to determine Aβ amino-acid sequence. Brain tissues were frozen or stored in RNAlater (Thermo Fisher Scientific, USA). To extract total RNA, 1 mL of ISOGEN (NIPPON GENE, Japan) was added to 50 μg of brain tissue, emulsified with Biomasher II (Nippi, Japan), then centrifuged at 12,000 ×*g* and 4°C for 15 min. Next, 500 μL of isopropanol was added to the aqueous layer, and the mixture was again centrifuged at 12,000 ×*g* and 4°C for 10 min. After the addition of 75% ethanol (500 µL), the precipitate was centrifuged a third time at 7,500 ×*g* and 4°C for 5 min. The supernatant was then air-dried for 10 min at room temperature (20-25°C). The final pellet was dissolved with 30 μL of Rnase-DNase free water to obtain total RNA for cDNA synthesis. Ten microliters of RNA were reverse-transcribed to cDNA using the OneStep RT-PCR kit (TAKARA). Primers (forward: 5’-GACTCACCACACGACCAGG-3’ and reverse: 5’-TCTTCTTCAGCATCACCAAAGT-3’) for PCR amplification of the Aβ region were developed based on budgerigar APP gene sequences. Thermocycling conditions (in a basic gradient thermocycler) were as follows: 35 cycles of 50°C for 30 min, 94°C for 2 min, 94°C for 30 s, 63°C for 30 s, and 72°C for 1 min. After thermocycling, either ExoSAP-IT (Affymetrix, USA) or Nucleo Spin Gel and PCR clean-up (TAKARA BIO, Japan) was added to purify amplicons. Sequencing was performed on the 3130 Genetic Analyzer (Applied Biosystems, USA).

A BLAST (http://blast.ncbi.nlm.nih.gov/Blast.cgi) was performed to obtain homologous sequences, which were analyzed in MEGA6 [9]. Nucleotide sequences were then converted into amino acids, and substitutions were identified. Furthermore, the APP gene sequences of all birds in NCBI were searched for their Aβ regions, which were then aligned with the human sequence to perform a phylogenetic analysis.

#### 5. Mass spectrometric analysis

Amyloid proteins were extracted from tissue and were digested with trypsin as described [10]. Digests were placed in a packed nano-capillary column (NTCC-360/75-3-123; 0.075 mm I.D. × 125 mm L, particle diameter 3 µm, Nikkyo Technos Co., Ltd., Tokyo, Japan) for separation in a DiNa HPLC system fitted with an automatic sampler (KYA Technology Corporation, Tokyo, Japan). The flow rate was 200 nL/min with a 2–80% linear gradient of acetonitrile in 0.1% formic acid. Eluted peptides were directly detected with an ion trap mass spectrometer, Velos Pro (Thermo Fisher Scientific Inc., Waltham, USA). Spectra were analyzed with Proteome Discoverer (Thermo Fisher Scientific Inc., Waltham, USA) and Mascot software (Matrix Science Inc., Boston, USA).

## Results

Of the 28 birds examined, only one (30–40 years old female Amazon parrot) exhibited CAA and slight senile plaque-like Aβ deposits. Prior to euthanasia, the female had already experienced two weeks of neurological disease before being found unmoving at the cage bottom. We were able to observe an 8 mm diameter blood clot in the cerebral ventricle of the right hemisphere, with evidence of bleeding—accompanied by encephalomalacia—in the surrounding brain tissue (Figs. 1 and 2). Histologically, CAA was present on the cerebral surface and deeper down (Fig. 3), diffusely deposited in small arterial walls of the cortex and meninges; indeed, CAA was so advanced that amyloid had replaced the walls (Fig. 4). Amyloid had a slight tendency to radially deposit in extravascular regions, scattered around blood vessels. When stained with Congo red, deposits emitted green polarized light (Fig. 4). Dual fluorescent immunostaining of anti-Aβ1-40 and 1-42 antibodies revealed orthotopic deposition in vessel walls (Fig. 5). The cerebral cortex contained several PAM-positive lesions similar to typical plaques, but Congo red staining was negative (Fig. 6). In addition, CAA was not deposited in cerebral cortex capillaries and we did not observe hyperphosphorylated tau accumulation. Under Kluver-Barrera staining, we saw myelin sheath degeneration within encephalomalacia foci, accompanied by macrophage infiltration. Immunostaining with anti-GFAP antibody revealed an increase in activated astrocytes around blood vessels. Holzer staining then identified fibrous astrocytes under the cerebral meninges, but no subcutaneous bleeding or lesions beyond CAA, known to cause cerebral hemorrhaging.

**Fig 1.**
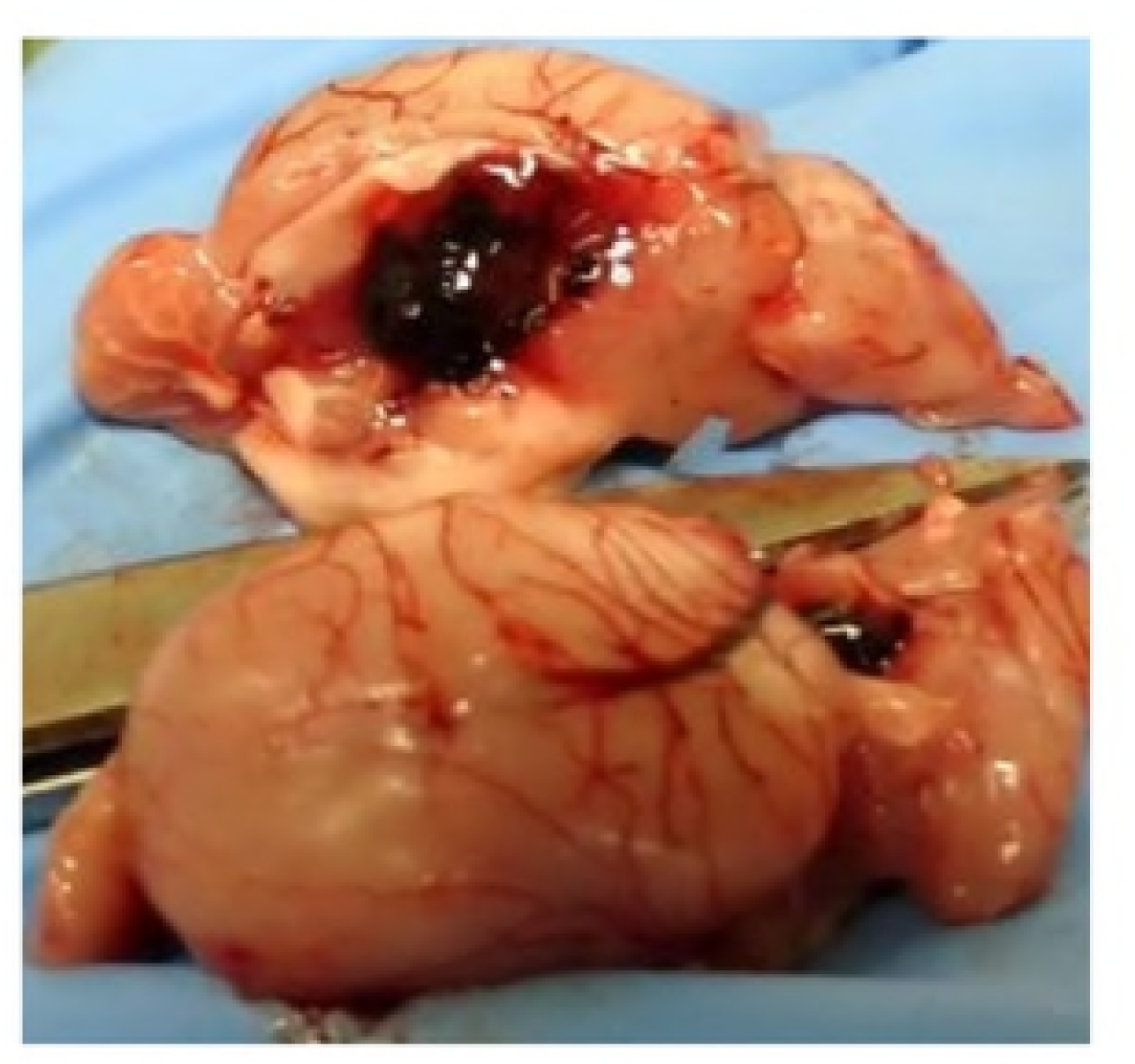
Brain of Amazon parrot showing 8 mm diameter blood clot

**Fig 2.**
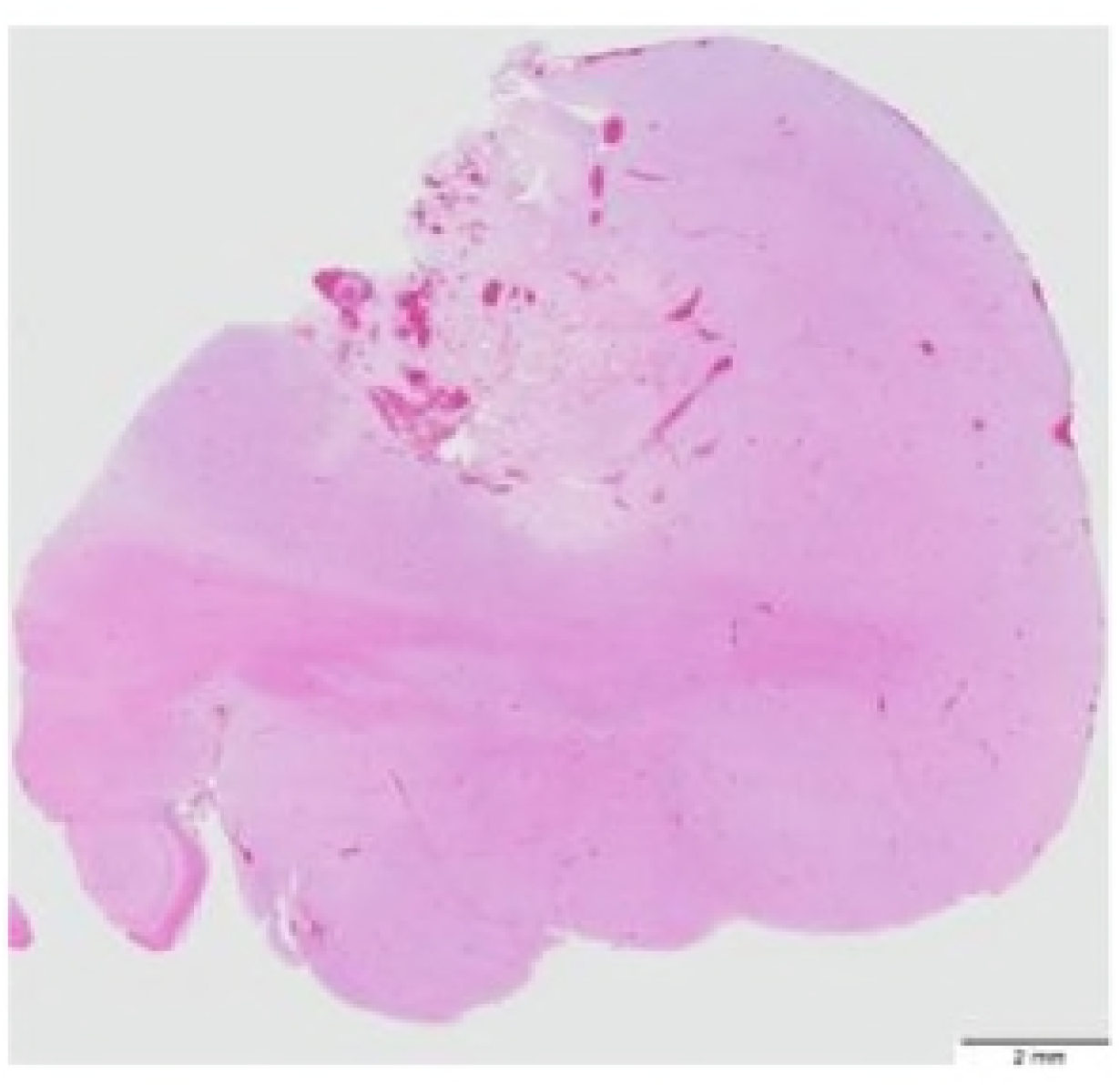
Multiple hemorrhages accompanied by cerebral encephalomalacia in Amazon parrot, visualized through H & E-staining and a magnifying loupe.

**Fig 3.**
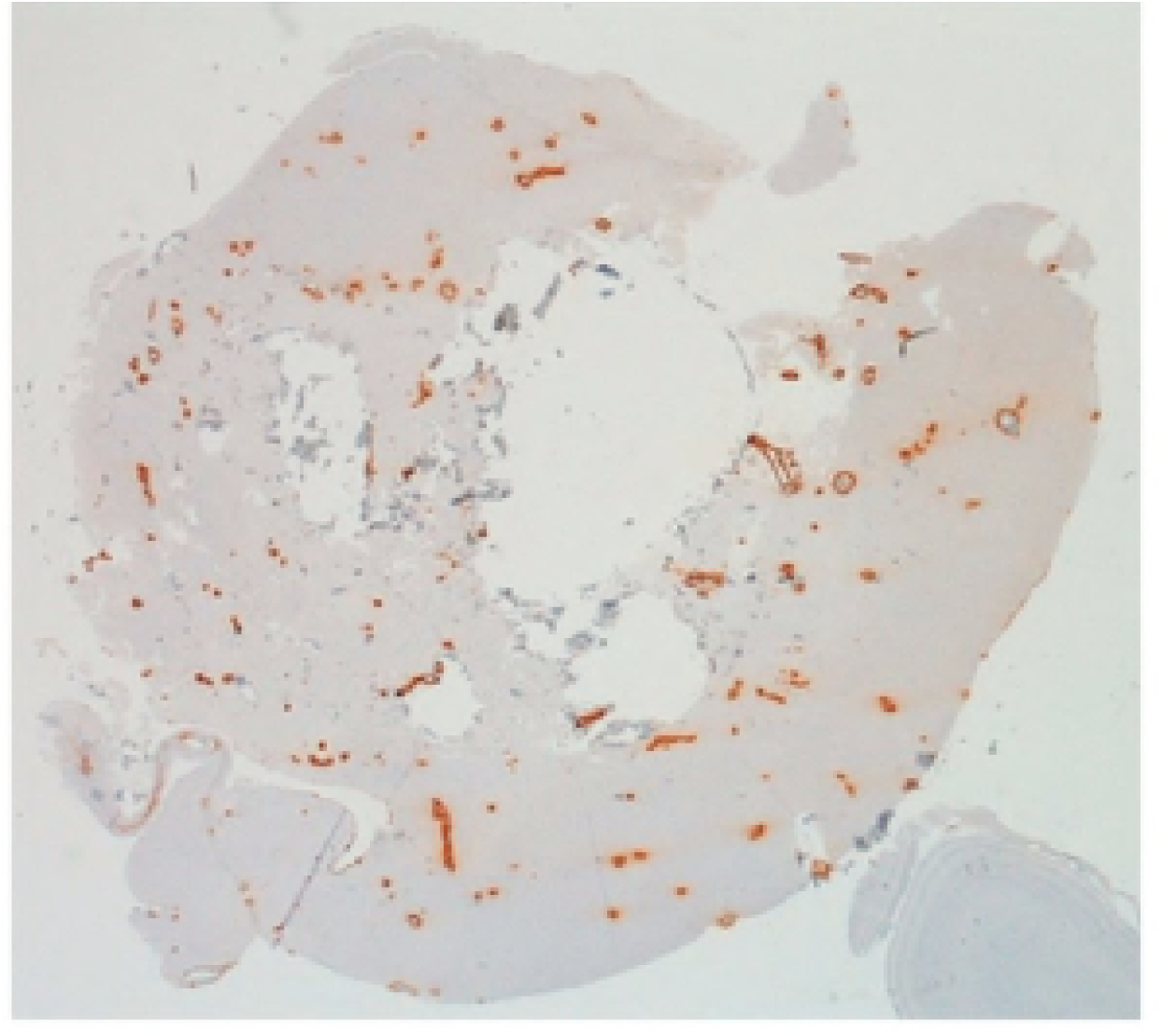
Cerebrum of Amazon parrot. Immunostaining with anti-Aβ 1-40 antibody revealed that CAA was widely distributed in the cerebral surface and cortex.

**Fig 4.**
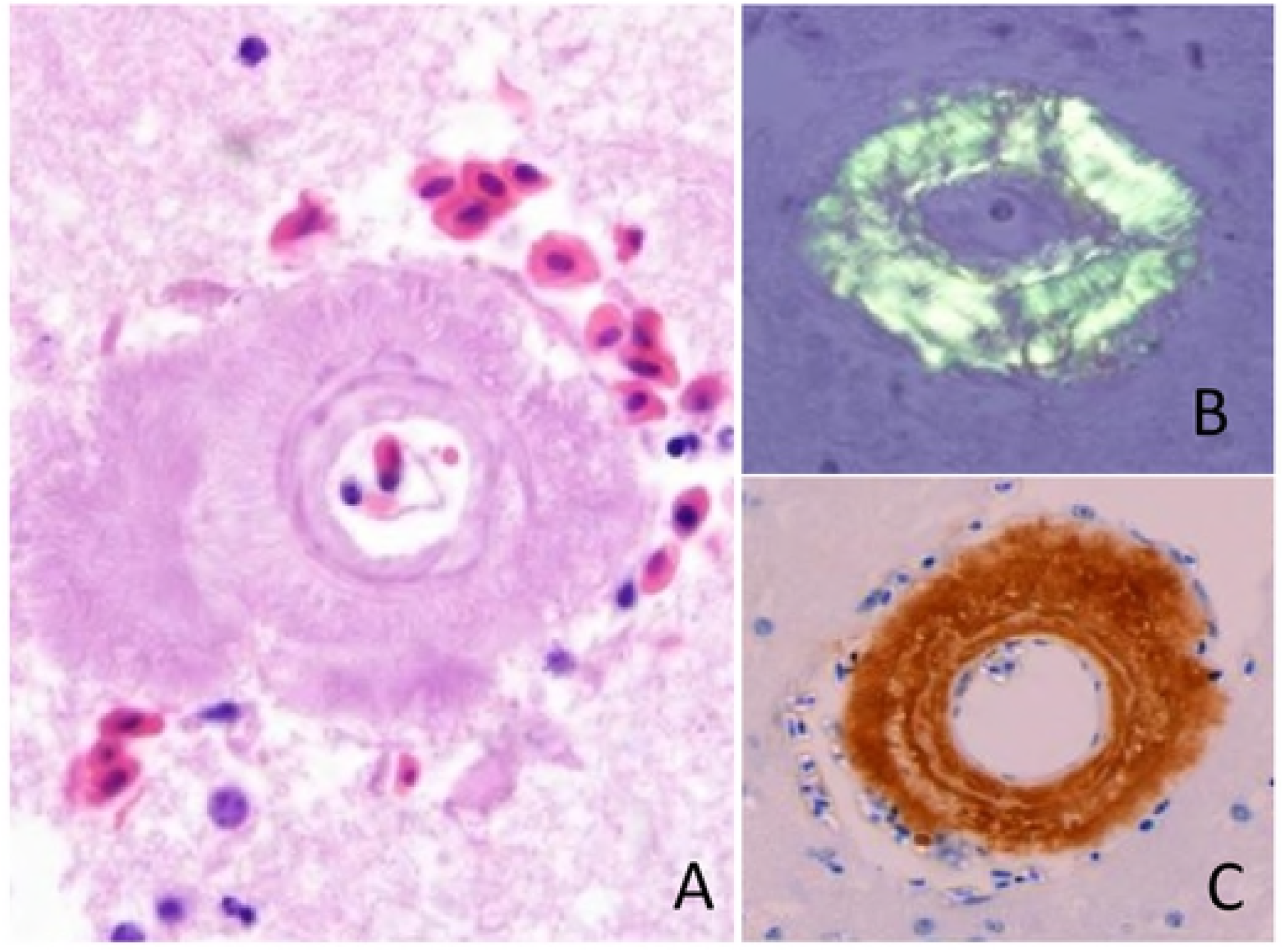
Cerebrum of Amazon parrot. (A) High magnification of vascular lesion, severe Aβ deposition, H & E-staining. (B) Congo Red stained tissue exhibiting green polarized light. (C) Immunostaining with anti-Aβ1-42 antibody revealed deposited Aβ.

**Fig 5.** The cerebrum of Amazon parrot. (A) anti Aβ 1-42 antibody, (B) anti Aβ 1-40 antibody, (C) merged image of double immunostaining.

**Fig 6.** Cerebrum of Amazon parrot showing senile plagues similar to typical plaques found in the cerebral cortex, revealed by PAM staining.

Amplicons of the target length (∼250 bp) were obtained in all nine samples after reverse-transcription PCR and the entire Aβ region successfully sequenced. This sequence was then aligned with homologous sequences obtained from NCBI. We identified a slight difference in nucleotide sequence across all seven species, but none resulted in amino acid substitution; thus, every sequence was consistent with human Aβ.

We then examined the Aβ region of 47 avian species and split them into three types based on amino-acid sequences: normal human (Type 1), valine (V) substitution of human type 31-isoleucine (I) (Type 2), and lysine (K) substitution of human type 3-glutamic acid (E) (Type 3). *Geospiza fortis, Serinus canaria*, and *Zonotrichia albicollis* Aβ sequences were all Type 2, while *Phaethon lepturus* was Type 3. The remaining species were classified as Type 1 (Table 2). Regardless of the presence or absence of Aβ deposition, predicted Aβ amino-acid sequences were Type 1 in all 28 birds that were subjected to pathological analyses.

Finally, through mass spectrometry of the Aβ protein isolated from the Amazon parrot sample, we identified a peptide corresponding to Aβ 6-16, 17-28, and 36-42 (Fig. 7). Although we did not detect Aβ 1-5 and 29-35, the amino-acid sequence of the deposited protein revealed was consistent with human Aβ.

**Fig 7.** Spectrum of peptides identified via LC-MSM. Mass spectrometry analysis of the Aβ protein deposited in the Amazon parrot cerebrum detected a peptide corresponding to Aβ 6-16, 17-28, and 36-42.

## Discussion

Aβ deposition in the brain can occur extracellularly or as CAA. In humans afflicted with Alzheimer’s disease, a major lesion is extracellular Aβ deposition, which can form senile plagues (Alzheimer A, 1911) or CAA, often simultaneously [11]. However, Aβ deposition modes vary across species, with CAA being common more in dogs [12], for example, but less common in cheetahs [13]. Here, we observed CAA in one sample, corresponding to past reports[7, 8], suggesting that this deposition mode is more likely in birds. Clinicopathological signs of CAA in humans include cerebral hemorrhage, ischemic lesions, and dementia [11, 14]. In dogs, amyloid deposits in capillaries, along with small and medium arteries, occur with increasing age. These deposits narrow the intravascular lumen and weaken blood-vessel walls, eventually causing ischemic lesions and cerebral hemorrhages [15]. In this study, we confirmed neurological symptoms in the sole case of CAA observed, and through pathological examination, diagnosed the cause of death as severe cerebral hemorrhage. This is the first clinical evidence of avian CAA.

In humans, old age is the most important risk factor for CAA [11], similar to what has been reported for other mammals [6]. Our study suggests that age is also a risk factor for avian CAA, because birds with this lesion were all older than their average life expectancy. Thus, Aβ deposition in avian species may begin once an individual survives beyond the average lifespan. These data imply the need for estimating animal life span, and several methods do exist to serve that purpose, including Rubner’s law of body surface [16]. Avian lifespan is very long compared with mammals of similar weight [17]. However, we should note that seven Humboldt penguins did not exhibit Aβ deposition despite being the same age as those that did, suggesting clear interspecific differences in the manifestation of this symptom.

In Alzheimer’s patients, CAA is thought to stem from Aβ 42 production in neurons that then are deposited in blood vessels, while Aβ 40 continues to deposit [11]. On the other hand, Shinkai et al. reported that Aβ 40 with low aggregability tends to deposit in blood vessels [18]. This study identified two types of orthotopic deposits in birds, but we are unable to discuss how Aβ type affects deposition susceptibility because no previous report has produced similar detailed data in birds. However, previous descriptive studies suggested that Aβ is deposited radially in the vessel wall outward [7, 8]. In contrast, our study found only a weak tendency to radial deposition; instead, we saw a satellite pattern around deposited blood vessels.

Deposition differences may be related to Aβ amino-acid sequences [6,19]. In birds, Aβ has at least three types of amino-acid sequences, but Type 1 was the most common. Indeed, Type 1 was found for the Amazon parrot and old Humboldt penguins without Aβ deposits in our study, along with birds examined in previous research [7,8]. These results suggest that CAA occurrence and morphology did not involve differences in amino-acid sequences. Instead, protein conformation and the influence of other aggregation factors were the main source of CAA variation. Furthermore, as mammalian Aβ sequences tended to be Type 1 (except in cats), the Aβ sequence appears to be well preserved, with high homology among homeothermic animals.

In conclusion, our results provide an important first step to accurately grasp Aβ deposition in birds. Given the importance of age, future studies should aim to sample from very old animals when investigating Aβ deposition in avian brains. Aging is a clear risk factor for Aβ deposition. Given avian longevity and the highly conserved nature of APP across animals, we suggest that birds may be a suitable model for studying the mechanism underlying Aβ deposits. In turn, such data should contribute to improving our knowledge of pathogenesis of Alzheimer’s disease and aid in therapeutic development.

## Acknowledgments

We thank the various zoos and institutions for providing bird specimens. We also would like to thank Editage (www.editage.jp) for English language editing. This study was supported by MEXT*-Supproted Program for the Private University Resesarch Branding Project,2016-2020 (Ministry of Education,Culture,Sports,Science and Technology)

